## Supporting Information for "Identification of novel plasma proteomic biomarkers of Dupuytren Disease"

| **S1 Table. 54 Differentially expressed genes in the SomaScan Hypothesis-free analysis.** | | | | | | |
| --- | --- | --- | --- | --- | --- | --- |
| **Gene** | **Express** | **p-val** | **FDR** | **Nodes** | **Notes** | **Cat** |
| *ACAN** | Up | 0.001148 | 0.218 | 2 | Aggrecan core protein 2; Proteoglycan, binds to hyaluronic acid. ***ACAN is near DD-related SNP rs6496519 and is dysregulated in DD transcriptomic profiling*.** *Extracellular matrix interactions are key components of DD biology.* [55, 59, 78] | 2, 4 |
| *AKR1A1* | Down | 0.001021 | 0.218 | 1 | Aldo-keto reductase family 1 member A1. *A potential DD relationship is unclear*. | ? |
| *AOC3** | Up | 0.000845 | 0.218 | 0 | Membrane primary amine oxidase; cell adhesion protein. **Increased *AOC3 levels in small vessel endothelium cells in DD-affected tissues*.** (referred to by alias *VAP-1*), *DD is associated with local microvascular inflammation, thrombosis, and endothelial leucocyte adhesion.* [13, 79] | 7 |
| *ARG2* | Up | 0.000281 | 0.211 | 0 | Arginase-2, mitochondrial; promotes endothelial senescence, monocyte adhesion, and enhances *VCAM1 ⁄ ICAM1* levels. *DD is associated with local microvascular inflammation, thrombosis, and endothelial leucocyte adhesion*. [13, 80] | 7 |
| *BIN2* | Down | 0.001793 | 0.246 | 0 | Bridging integrator 2; Promotes cell motility and migration; central regulator of platelet activation in thrombosis and thrombo-inflammatory disease settings. *Cell-matrix, cell-cell, and cytoskeletal interactions are core processes in DD. DD is associated with local microvascular inflammation, thrombosis, and endothelial leucocyte adhesion.* [13, 78, 81] | 5, 7 |
| *C5* | Down | 0.001610 | 0.225 | 3 | Complement C5 alpha chain; initiates complement components, C5-C9 resulting in the Terminal Complement Complex, which is increased in the serum of Idiopathic Pulmonary Fibrosis; activates basophil and mast cell degranulation responsible for urticaria. *Itching is common in DD-affected tissues.* [82-84] | 3 |
| *C6* | Down | 0.001610 | 0.225 | 1 | Complement component C6. Forms part of the Terminal Complement Complex which is increased in the serum of Idiopathic Pulmonary Fibrosis. Upregulated in healing tendons subjected to mechanical loading; *DD fibroblast gene expression responds to mechanical loading.* [84-86] | 5 |
| *CALCB* | Up | 0.000333 | 0.211 | 1 | Calcitonin gene-related peptide 2. *A potential DD relationship is unclear.* | ? |
| *CASP3** | Down | 0.001603 | 0.075 | 8 | Caspase-3 subunit p12; Involved in apoptosis. ***PPI Network analysis projects key involvement in DD***. *Apoptosis is dysregulated in DD*. ***CASP3 is dysregulated in DD transcriptomic profiling***. [55, 87, 88] | 1, 2 |
| *CGA* | Up | 0.001093 | 0.218 | 2 | Glycoprotein hormones alpha chain; expression increases with age; *DD prevalence increases with age*. ***CGA is dysregulated in DD transcriptomic profiling***. [55, 82, 89] | 3 |
| *CGB3* | Up | 0.001093 | 0.218 | 1 | Chorionic gonadotropin subunit beta 3. *A potential DD relationship is unclear*. | ? |
| *CGB7* | Up | 0.001093 | 0.218 | 0 | Choriogonadotropin subunit beta 7. *A potential DD relationship is unclear*. | ? |
| *CPB1* | Up | 0.001398 | 0.222 | 2 | Carboxypeptidase B1. *A potential DD relationship is unclear*. | ? |
| *CRKL* | Down | 0.000892 | 0.218 | 3 | Crk-like protein; needed for fibroblast cytoskeletal structure and motility; epigenetic regulation of fibroblast focal adhesions. *Cell-matrix, cell-cell, and cytoskeletal interactions are core processes in DD*. [78, 90, 91] | 5 |
| *CSNK1G2** | Down | 0.001222 | 0.218 | 0 | Casein kinase I isoform gamma-2; Serine/threonine-protein kinase. Participates in WNT signaling. *WNT expression is dysregulated in DD; dysregulated in DD transcriptomic profiling.* ***Pathway analysis projects differential CSNK1G2 expression in both visibly affected and normal-appearing tissues in DD vs. control***. [55, 92, 93] | 2 |
| *DDX19A* | Down | 0.000886 | 0.218 | 0 | ATP-dependent RNA helicase DDX19A. *A potential DD relationship is unclear*. | ? |
| *DSG2* | Up | 0.001607 | 0.225 | 1 | Desmoglein-2; Fundamental component of intercellular desmosome junctions and cell-cell adhesion; involved in mechanosensitive and mechanoresponsive pathways. *Cell-matrix, cell-cell, and cytoskeletal interactions are core processes in DD*. [78, 94, 95] | 5 |
| *EDIL3* | Up | 0.000921 | 0.218 | 0 | EGF-like repeat and discoidin I-like domain-containing protein 3; Promotes adhesion of endothelial cells through interaction with the alpha-v/beta-3 integrin receptor. *DD is associated with local microvascular inflammation, thrombosis, and endothelial leucocyte adhesion*. [13, 96] | 7 |
| *EIF4H* | Down | 0.001031 | 0.218 | 1 | Eukaryotic translation initiation factor 4H; Stimulates the RNA helicase activity of EIF4A in the translation initiation complex. *EIF4A and ELN genes are in close proximity, and elastin loss is characteristic of DD-affected tissues*. [97, 98] | 4 |
| *ELAPOR1* | Up | 0.000609 | 0.218 | 0 | Endosome/Lysosome-Associated Apoptosis And Autophagy Regulator 1; involved in apoptosis and cell proliferation; may protect cells from cell death (referred to by alias EIG121). *Apoptosis is dysregulated in DD*. [88, 99, 100] | 1 |
| *FAM234B* | Up | 0.001227 | 0.218 | 2 | Protein FAM234B; predicted to be located in Golgi apparatus, cytoskeleton, and cell membrane. *Cell-matrix, cell-cell, and cytoskeletal interactions are core processes in DD*. [78] | 5 |
| *FKBP5* | Down | 0.001045 | 0.218 | 1 | Peptidyl-prolyl cis-trans isomerase FKBP5; involved in procollagen I triple helix assembly; *local collagen I accumulation is a prominent feature of DD. R*esponsible for the effects of mechanical loading on *COL1A1* and *COL1A2* expression (referred to by alias *PPIASE*); *DD fibroblast gene expression responds to mechanical loading*. [85, 101] | 4, 6 |
| *G3BP2* | Down | 0.001539 | 0.225 | 3 | Ras GTPase-activating protein-binding protein 2; binding varies with extracellular matrix stiffness; expression increased by mechanical stress; involved in endothelial shear stress-induced loss of endothelial barrier function, monocyte adhesion to endothelial cells, and proinflammatory cytokines. *DD is associated with local microvascular inflammation, thrombosis, and endothelial leucocyte adhesion. Cell-matrix, cell-cell, and cytoskeletal interactions are core processes in DD*. [13, 78, 102, 103] | 5, 7 |
| *GP1BB* | Down | 0.000767 | 0.218 | 1 | Platelet glycoprotein Ib beta chain; transmembrane platelet protein; binds to subendothelial von Willebrand factor; involved in forming platelet plugs; necessary for normal platelet adhesion to a stimulated endothelium. *DD is associated with local microvascular inflammation, thrombosis, and endothelial leucocyte adhesion*. [13, 104] | 7 |
| *GUSB* | Down | 0.000113 | 0.185 | 1 | Beta-glucuronidase; degrades extracellular matrix glycosaminoglycans, including heparan sulfate, dermatan sulfate, and chondroitin-4,6-sulfate. *Extracellular matrix remodeling is a prominent process in DD*. [78, 105] | 5 |
| *HGS* | Down | 0.000178 | 0.208 | 7 | Hepatocyte growth factor-regulated tyrosine kinase substrate. *A potential DD relationship is unclear*. | ? |
| *HSP90AA1* | Down | 0.001139 | 0.218 | 0 | Heat shock protein HSP 90-alpha; activates endothelial (KNG1) bradykinin-forming cascade; activates MMP2; in combination, TGF-β1 and Hsp90β stimulate anchorage-independent growth, reduce adhesion and stimulate migration via an alternate TGF-β1 pathway, mediated by αvβ6 integrin. *Cell-matrix, cell-cell, and cytoskeletal interactions are core processes in DD*. [78, 106, 107] | 4, 5 |
| *KNG1** | Up | 0.000490 | 0.218 | 6 | Kininogen1; Alternative splicing produces high molecular weight kininogen (HMWK) and low molecular weight kininogen (LMWK). HMWK inhibits thrombin- and plasmin-induced thrombocyte aggregation; stimulates the release of other mediators of inflammation; causes vasodilation and increases vascular permeability; the bradykinin B2 receptor contributes to endothelial inflammation. *DD is associated with local microvascular inflammation, thrombosis, and endothelial leucocyte adhesion*. [13, 108] | 7 |
| *MAP3K11* | Up | 0.001532 | 0.225 | 2 | Mitogen-activated protein kinase kinase kinase 11; plays a role in the activation of BRAF. *BRAF inhibitors provoke DD-like clinical changes*. [109] | 3 |
| *MBNL1* | Down | 0.001278 | 0.218 | 1 | Muscleblind-like protein 1; central regulator of fibrosis; regulates αSMA; regulates the TGF-β/Smad signaling pathway; *SMAD - TGF are prominent pathways in DD fibroblast proliferation and contraction*. [57, 110] | 6 |
| *MBNL2* | Down | 0.000971 | 0.218 | 1 | Muscleblind-like protein 2; Mediates pre-mRNA alternative splicing regulation; part of an RNA-binding protein network governing fibroblast to myofibroblast transition. *Fibroblast differentiation into myofibroblast phenotype is a core process in DD*. [111, 112] | 6 |
| *NAA80* | Down | 0.001251 | 0.218 | 0 | N-alpha-acetyltransferase 80; regulates actin filament acetylation, depolymerization, and elongation; plays essential roles in filament assembly, cytoskeleton organization, and cell motility. *Cell-matrix, cell-cell, and cytoskeletal interactions are core processes in DD*. [78, 113] | 5 |
| *NRG4* | Up | 0.000669 | 0.218 | 1 | Pro-neuregulin-4, membrane-bound isoform; brown fat adipokine; lower levels in obese compared to normal-weight. *DD is associated with loss of subcutaneous palm fat and lower triceps skinfold thickness compared with normal*. [114-116] | 3 |
| *OSCAR* | Up | 0.000070 | 0.185 | 1 | Osteoclast-associated immunoglobulin-like receptor; collagen I is an *OSCAR* ligand. *OSCAR* also stimulates the release of TNFα from T-cells. *TNF is involved in DD pathways*. [61, 66] | 2 |
| *PCSK9* | Up | 0.001371 | 0.222 | 1 | Proprotein convertase subtilisin/kexin type 9; blocks LDL degradation and increases circulating LDL. *DD is associated with increased mortality from cardiovascular disease*. [71] | 3 |
| *POSTN** | Up | 0.000312 | 0.211 | 2 | Periostin; Induces cell attachment and spreading and plays a role in cell adhesion. Enhances incorporation of BMP1 in the fibronectin matrix of connective tissues, and subsequent proteolytic activation of lysyl oxidase LOX. *Increased LOX activity in Dupuytren tissue.* ***Increased POSTN expression in DD fibroblasts and in sweat glands of DD-adjacent skin****.* [55, 94, 117-119] | 4, 6 |
| *PRKAR2B* | Down | 0.001119 | 0.218 | 0 | cAMP-dependent protein kinase type II-beta regulatory subunit. *A potential DD relationship is unclear*. | ? |
| *PRSS1* | Up | 0.000809 | 0.218 | 1 | Alpha-trypsin chain 1; Belongs to the peptidase S1 family; expressed in the pancreas and some malignancies. Activates several metalloproteinases, but only where locally expressed. *A potential DD relationship is unclear*. | ? |
| *RAB24* | Down | 0.000235 | 0.211 | 0 | Ras-related protein Rab-24; involved in autophagy-related processes; overexpressed in senescent fibroblasts. *DD fibroblasts have altered senescence pathways*. [120] | 1 |
| *RHOT1* | Down | 0.000133 | 0.185 | 0 | Mitochondrial Rho GTPase 1; essential for dynamic positioning of mitochondria along the length of axons; mitochondrial positioning along the length of axons and trafficking; impaired function is associated with neurodegenerative disease; (referred to by alias MIRO). *Progressive Pacinian corpuscle denervation is associated with DD*. [121, 122] | 3 |
| *SCGN* | Up | 0.000085 | 0.185 | 0 | Secretagogin, cytoplasmic calcium binding protein. *A potential DD relationship is unclear*. | ? |
| *SCPEP1* | Down | 0.000439 | 0.218 | 0 | Retinoid-inducible serine carboxypeptidase*. A potential DD relationship is unclear*. | ? |
| *SERPINC1* | Up | 0.001500 | 0.225 | 4 | Antithrombin-III; regulates the blood coagulation cascade; inhibits thrombin, and factors IXa, Xa and Xia; binds to endothelial heparin-like molecules. Deficiency or loss-of-function variants are major risk factors for thromboembolic disease. (referred to by alias Antithrombin III). *DD is associated with local microvascular inflammation, thrombosis, and endothelial leucocyte adhesion*. [13] | 7 |
| *SHMT1* | Down | 0.000619 | 0.218 | 0 | Serine hydroxymethyltransferase, cytosolic; Interconversion of serine and glycine. Nearly one third of collagen fibril composition is glycine, but the *potential impact of SHMT1 on DD-related collagen accumulation is unclear*. | ? |
| *SMAD1** | Down | 0.001275 | 0.218 | 0 | Mothers against decapentaplegic homolog 1; Transcriptional modulator activated by BMP (bone morphogenetic proteins) type 1 receptor kinase. SMAD1/5/9 pathway action is antifibrotic in multiple organ fibroses. *SMAD-TGFβ pathways are prominent in DD*. ***SMAD1 protein expression is normal in DD tissues, but intracellular SMAD1 mRNA expression is reduced in DD fibroblasts*.** [57] | 2, 6 |
| *SOCS3* | Down | 0.001007 | 0.218 | 3 | Suppressor of cytokine signaling 3. SOCS3 is involved in negative regulation of cytokines that signal through the JAK/STAT pathway. *Feedback loop regulation of DD-related cytokines, including IL6 and TNF*. *DD-related fibrosis involves JAK/STAT/IL13 signaling pathways*. [123] | 2 |
| *SPART* | Down | 0.000247 | 0.211 | 0 | Spartin; Participates in cytokinesis. SPART is associated with Epidermal Growth Factor Receptor (EGFR) degradation and transport to the cell membrane; *an increased ratio of intracellular to cell membrane EGFR has been found in early DD disease and is upregulated in cell culture.* ***SPART is under-expressed in DD Fibroblasts by Genome-wide exon expression profiles*** (referred to by alias SPG20). [58, 68] | 2, 6 |
| *SPATA22* | Up | 0.000871 | 0.218 | 0 | Spermatogenesis-associated protein 22. *A potential DD relationship is unclear*. | ? |
| *SYK* | Down | 0.000513 | 0.218 | 6 | Tyrosine-protein kinase SYK; regulates focal adhesion kinase. Integrins and related cell membrane adhesion and mechanosensing domains. *Cell-matrix, cell-cell, and cytoskeletal interactions are core processes in DD*. [78, 124] | 5 |
| *TBC1D13* | Down | 0.001076 | 0.218 | 0 | TBC1 domain family member 13. *A potential DD relationship is unclear*. | ? |
| *TF** | Up | 0.000025 | 0.172 | 2 | Serotransferrin; Responsible for the transport of iron from sites of absorption and heme degradation; iron levels are increased in the lung tissues of patients with idiopathic pulmonary fibrosis; exogenous iron increases human lung fibroblast proliferation and cytokine responses. *Local tissue iron deposition occurs in the early cellular stage of DD*. [125, 126] | 4, 7 |
| *USP8** | Down | 0.001398 | 0.222 | 2 | Ubiquitin carboxyl-terminal hydrolase 8; Hydrolase that can remove conjugated ubiquitin; prevents Wnt receptor frizzled 5 (FZD5); USP8 pathway is essential for Wnt/β-catenin signaling; *WNT signaling pathways are central in DD biology*. ***USP8 is under-expressed in DD Fibroblasts by Genome-wide exon expression profiles***. USP8 may exert protective influences against aging; *DD prevalence is age-related*. [10, 58] | 2, 3 |
| *WFIKKN2* | Up | 0.001128 | 0.218 | 0 | WAP, Kazal, immunoglobulin, Kunitz, and NTR domain-containing protein 2; Probably has serine protease- and metalloprotease-inhibitor activity. *MMP inhibition plays a role in DD biology*. Regulates the balance between the activation of Smad and non-Smad pathways by TGFB1. *SMAD-TGFβ pathways are prominent in DD.* [57] | 2, 6 |
| *YWHAB* | Down | 0.001257 | 0.218 | 4 | 14-3-3 protein beta/alpha, N-terminally processed. *A potential DD relationship is unclear*. | ? |

| **S2 Table. Functionally enriched pathways in the hypothesis-free analysis.** | | | | | |
| --- | --- | --- | --- | --- | --- |
| **Category** | **ID** | **Description** | **STR** | **FDR** | **Genes of matching proteins in this network** |
| GO Function | GO:0005102 | Signaling receptor binding | 0.59 | 0.0084 | *C5, EDIL3, GUSB, PCSK9, CASP3, WFIKKN2, HSP90AA1, CRKL, CGB3, SYK, NRG4, TF, CALCB, CGB7, CGA, KNG1* |
| GO Function | GO:0004857 | Enzyme inhibitor activity | 0.87 | 0.0322 | *C5, PRKAR2B, CASP3, WFIKKN2, SOCS3, SERPINC1, YWHAB, KNG1* |
| GO Function | GO:0045309 | Protein phosphorylated amino acid binding | 1.41 | 0.0381 | *SOCS3, CRKL, YWHAB, SYK* |
| GO Component | GO:0005615 | Extracellular space | 0.5 | 5.52E-06 | *C5, DSG2, SCPEP1, C6, PRKAR2B, EDIL3, GUSB, PCSK9, PRSS1, WFIKKN2, SHMT1, HGS, HSP90AA1, CGB3, SERPINC1, KIAA1324, AKR1A1, YWHAB, POSTN, NRG4, TF, CPB1, CALCB, FKBP5, CGB7, OSCAR, CGA, KNG1* |
| GO Component | GO:0005576 | Extracellular region | 0.43 | 7.90E-06 | *C5, DSG2, SCPEP1, C6, PRKAR2B, EDIL3, GUSB, PCSK9, PRSS1, WFIKKN2, SHMT1, HGS, HSP90AA1, CGB3, SERPINC1, KIAA1324, AKR1A1, YWHAB, SCGN, POSTN, NRG4, TF, ACAN, CPB1, CALCB, FKBP5, CGB7, OSCAR, CGA, KNG1, BIN2* |
| GO Component | GO:0031982 | Vesicle | 0.38 | 0.0025 | *C5, DSG2, SCPEP1, C6, PRKAR2B, EDIL3, GUSB, PCSK9, RAB24, AOC3, SHMT1, HGS, HSP90AA1, SERPINC1, KIAA1324, AKR1A1, YWHAB, SYK, SCGN, USP8, TF, CPB1, FKBP5, OSCAR, KNG1, BIN2* |
| GO Component | GO:0070062 | Extracellular exosome | 0.5 | 0.0035 | *C5, DSG2, SCPEP1, C6, PRKAR2B, EDIL3, GUSB, SHMT1, HGS, HSP90AA1, SERPINC1, KIAA1324, AKR1A1, YWHAB, TF, FKBP5, OSCAR, KNG1* |
| GO Component | GO:0019897 | Extrinsic component of plasma membrane | 1.02 | 0.0327 | *PCSK9, CRKL, SYK, USP8, TF* |
| GO Component | GO:0062023 | Collagen-containing extracellular matrix | 0.8 | 0.0327 | *EDIL3, PRSS1, HSP90AA1, SERPINC1, POSTN, ACAN, KNG1* |
| KEGG | hsa04610 | Complement and coagulation cascades | 1.25 | 0.0301 | *C5, C6, SERPINC1, KNG1* |
| Reactome | HSA-162582 | Signal Transduction | 0.46 | 0.0148 | *C5, CSNK1G2, DSG2, PRKAR2B, SMAD1, MAP3K11, CASP3, SOCS3, HGS, HSP90AA1, CRKL, RHOT1, YWHAB, SYK, NRG4, USP8, CALCB, FKBP5, CGA, KNG1* |

**S3 Table. 328 protein-coding genes in the SomaScan hypothesis-based analysis.**

*A1BG, A2M, ACAN, ACAT1, ACVR1B, ADAM12, ADAM15, ADAMTS3, ADH1B, ADK, AGA, AGT, AKR1C2, ALDH2, AMT, ANGPTL4, ANGPTL7, AOC3, APOB, ARHGEF10, ARHGEF2, ATXN3, AXIN2, AZGP1, B2M, BCL2, BDNF, BGN, BMP1, BMP6, BSG, C3, CADM1, CANT1, CARD18, CASP3, CASP8, CAT, CCL2, CCL5, CCN2, CCN4, CD34, CD4, CD40LG, CD44, CD68, CD8A, CDH1, CDH11, CDH13, CDH4, CDKN1A, CERT1, CFDP1, CFP, CHI3L1, CHRD, CHST6, CLU, CNTN2, CNTN6, COL10A1, COL11A2, COL13A1, COL15A1, COL18A1, COL1A1, COL20A1, COL23A1, COL25A1, COL28A1, COL2A1, COL3A1, COL5A1, COL6A1, COL6A2, COL6A3, COL6A5, COL8A1, COL9A1, COL9A3, COLGALT1, COLGALT2, COQ7, CSF2, CSMD1, CSNK1G2, CTHRC1, CTNNB1, CTSK, CXCL1, CXCL14, CXCL8, DAB2, DCN, DDR1, DDR2, DES, DKK1, DPP10, DPP4, DSCAML1, EED, EGF, EGFR, EGLN1, EGLN2, EGLN3, ELANE, ENG, EPDR1, EYS, F13A1, FAM20A, FBP1, FGF2, FGFR2, FKBP4, FMOD, FN1, FRZB, FST, G0S2, GCKR, GDF5, GH1, GJA1, GLB1, GPC1, GPR142, GPT, GPX3, GRIA4, HBEGF, HGF, HIF1A, HLA-C, HLA-G, HMGA2, HPRT1, HPX, HSD17B7, HSF1, HSPG2, ICAM1, IFNG, IGF1, IGF1R, IGF2, IGFBP6, IGFBP7, IKBKB, IL13, IL17A, IL1A, IL1B, IL6, INHBA, INS, INSR, IST1, ITGA11, ITGA2, ITGA4, ITGA5, ITGA6, ITGB1, KDR, KIN, KITLG, KNG1, KPNA2, KRT1, KRT34, LAMA3, LAMB1, LAMC2, LCN2, LOXL2, LOXL3, LRP5, LUM, LY75, MAP2K1, MAP4K5, MAPK3, MB, MECP2, MFAP5, MIF, MME, MMP1, MMP10, MMP12, MMP13, MMP14, MMP2, MMP20, MMP3, MMP7, MMP8, MMP9, MOCOS, MYC, NCALD, NCL, NDE1, NECAB1, NFKB1, NGF, NGFR, NID1, NME8, NOG, NRG1, NRXN1, NT5E, NTM, OSTF1, P3H1, P4HA1, P4HA2, P4HB, PCNA, PCOLCE, PCOLCE2, PCSK1N, PCSK7, PDGFA, PDLIM5, PECAM1, PIK3AP1, PLAT, PLCG1, PLG, PLOD2, PLOD3, PLXDC2, POLD4, POSTN, PPARA, PPIB, PRKCA, PROS1, PSTPIP1, PTGDS, PTGS2, PTK2, PTN, PTPRD, RAB31, RBFOX1, RND3, ROR2, RSPO2, S100A6, SERPINE1, SERPINH1, SFRP1, SFRP4, SH2D1A, SH3BP2, SH3GL2, SIRPB1, SLC14A2, SLMAP, SMAD1, SMAD2, SMAD3, SMAD4, SOD1, SOD2, SORCS3, SPARC, SPOCK3, SPP1, STAT1, STAT3, STATH, STIP1, STUB1, SUMO4, TAC1, TAGLN, TBK1, TCF4, TF, TFPI2, TGFB1, TGFB2, TGFB3, TGFBR1, TGFBR2, THBS2, THBS4, THY1, TIMP1, TIMP2, TIMP3, TMEM132A, TNC, TNF, TNFRSF1B, TNIP1, TNXB, TP53, TPI1, TXN, UBE2D3, UBE2K, USP8, UST, VAPA, VCAM1, VDR, VEGFA, VIM, VTN, WIF1, WNK1, WNT11, WNT3A, WNT5A, WWOX, YAP1, YWHAZ, ZNF264*

| **S4 Table. 23 Differentially expressed genes in the Hypothesis-based analysis.** | | | | | | |
| --- | --- | --- | --- | --- | --- | --- |
| **Gene** | **Exp** | **p-val** | **FDR** | **Nodes** | **Notes** | **Cat** |
| *ACAN** | Up | 0.001148 | 0.074 | 3 | Aggrecan core protein 2; Proteoglycan, binds to hyaluronic acid. ***ACAN is near DD-related SNP rs6496519 and is dysregulated in DD transcriptomic profiling***. Extracellular matrix interactions are key components of DD biology. [55, 59, 78] | 2, 4 |
| *AOC3** | Up | 0.000845 | 0.074 | 0 | Membrane primary amine oxidase; cell adhesion protein. Increased ***AOC3 levels in small vessel endothelium cells in DD-affected tissues***. (referred to by alias VAP-1), *DD is associated with local microvascular inflammation, thrombosis, and endothelial leucocyte adhesion.* [13, 79] | 7 |
| *CASP3** | Down | 0.001603 | 0.075 | 8 | Caspase-3 subunit p12; Involved in apoptosis. ***PPI Network analysis projects CASP3 involvement in DD****. Apoptosis is dysregulated in DD.* ***CASP3 is dysregulated in DD transcriptomic profiling***. [55, 87, 88] | 1, 2 |
| *COL1A1* | Up | 0.010705 | 0.211 | 7 | Collagen alpha-1(I) chain. *Homotrimer Collagen (a1) is increased in DD and resistant to enzymatic degradation. Differential expression in DD cell culture.* [31, 33, 55] | 4 |
| *CSMD1* | Up | 0.007311 | 0.191 | 0 | CUB and sushi domain-containing protein 1; Potential suppressor of squamous cell carcinomas; *CSMD1* variant associated with reduced severity of post-burn hypertrophic scarring. *Hypertrophic scar shares some ECM, microvascular, and mechanical load effects with DD*. [127-130] | 5, 7 |
| *CSNK1G2** | Down | 0.001222 | 0.074 | 0 | Casein kinase I isoform gamma-2; Serine/threonine-protein kinase. Participates in WNT signaling. WNT expression is dysregulated in DD; dysregulated in DD transcriptomic profiling. ***Pathway analysis projects differential CSNK1G2 expression in both visibly affected and normal-appearing tissues in DD vs. control***. [55, 92, 93] | 2 |
| *DAB2* | Down | 0.009245 | 0.211 | 1 | Disabled homolog 2; regulates cell membrane adhesion complex disassembly during cell migration. *Fibroblast adhesion and migration processes are abnormal in DD*. [131, 132] | 5 |
| *DDR2* | Up | 0.010921 | 0.211 | 3 | Discoidin domain-containing receptor 2; Tyrosine kinase that functions as a cell surface receptor for fibrillar collagen and regulates cell differentiation, remodeling of the extracellular matrix, cell migration, and cell proliferation. Regulates extracellular matrix remodeling by up-regulating the collagenases MMP1, MMP2, and MMP13. Promotes fibroblast migration and proliferation. *All of these processes are abnormal in DD.* ***DDR2 is near the DD-related SNP rs17433710***. [59, 78, 131] | 2,4,5 |
| *HPX* | Up | 0.013002 | 0.229 | 3 | Hemopexin; required for MMP1 activity. *MMP1 plays a key role in DD biology*. [37, 133] | 4 |
| *KNG1** | Up | 0.000490 | 0.069 | 3 | Kininogen1; Alternative splicing produces high molecular weight kininogen (HMWK) and low molecular weight kininogen (LMWK). HMWK inhibits thrombin- and plasmin-induced thrombocyte aggregation; stimulates the release of other mediators of inflammation; causes vasodilation and increases vascular permeability; the bradykinin B2 receptor contributes to endothelial inflammation. *DD is associated with local microvascular inflammation, thrombosis, and endothelial leucocyte adhesion*. [13, 108] | 7 |
| *LCN2* | Down | 0.001910 | 0.081 | 3 | LCN2 is an iron-ion binding protein; involved in multiple processes including innate immunity and ferroptosis. *Hemosiderin deposition occurs in the early cellular stage of DD*. LCN2 protects MMP9 from degradation. *MMP9 activates TGFB*. (referred to by alias LGA2) [84, 126, 134, 135] | 3, 4 |
| *MAP2K1* | Down | 0.003473 | 0.133 | 4 | Dual specificity mitogen-activated protein kinase kinase 1; MAP2K1 phosphorylation from mechanical stretch increases TGFB expression. Mechanical stretch of myofibroblasts attached to a stiff extracellular matrix activates latent TGFB1. *Increasing ECM stiffness increases smooth muscle actin expression and reduces migration of DD fibroblasts compared to control fibroblasts*. [131, 136, 137] | 4,5 |
| *PLAT* | Down | 0.006181 | 0.186 | 2 | Tissue-type plasminogen activator chain A; converts plasminogen to plasmin, which, in turn, activates TGFB. *TGFB1 and TGFB2 are involved in key DD-related pathways*. [138] | 6 |
| *POSTN** | Up | 0.000312 | 0.066 | 6 | Periostin; Induces cell attachment and spreading and plays a role in cell adhesion. Enhances incorporation of BMP1 in the fibronectin matrix of connective tissues, and subsequent proteolytic activation of lysyl oxidase LOX. *Increased LOX activity in Dupuytren tissue.* ***Increased POSTN expression in DD fibroblasts and in sweat glands of DD-adjacent skin***. [55, 94, 117-119] | 4, 6 |
| *PRKCA* | Down | 0.009837 | 0.211 | 4 | Protein kinase C alpha type; involved in the regulation of cell proliferation, apoptosis, differentiation, migration, and adhesion; multiple protein-protein interactions in DD network biology. (referred to by alias *PRKACA*). [139] | 1, 5 |
| *SERPINH1* | Down | 0.011476 | 0.211 | 3 | SERPINH1; collagen-binding protein that may affect the expression of collagen III. ***Increased SERPINH1 expression in DD tissue***. (referred to by alias *HSP47*). [140, 141] | 4, 5 |
| *SFRP4* | Down | 0.004311 | 0.152 | 2 | Secreted frizzled-related protein 4; inhibits WNT/frizzled receptor signaling pathway. ***Near DD-related SNPs rs16879765 and rs17171229.*** [64] | 2 |
| *SMAD1** | Down | 0.001275 | 0.074 | 1 | Mothers against decapentaplegic homolog 1; Transcriptional modulator activated by BMP (bone morphogenetic proteins) type 1 receptor kinase. SMAD1/5/9 pathway action is antifibrotic in multiple organ fibroses. ***SMAD1 protein expression is normal in DD tissues, but intracellular SMAD1 mRNA expression is reduced in DD fibroblasts.*** [57] | 2, 6 |
| *STAT1* | Down | 0.006707 | 0.189 | 4 | Signal transducer and activator of transcription 1-alpha/beta; mediates cellular responses to interferons and other cytokines and other growth factors; ***significantly reduced STAT1 expression in DD cell cultures.*** [58] | 6 |
| *STAT3* | Down | 0.005622 | 0.182 | 9 | Signal transducer and activator of transcription 3; Signal transducer and transcription activator that mediates cellular responses to interleukins and growth factors. ***STAT3 is near Dupuytren-associated risk alleles rs16879765 and rs17171229 risk alleles and is associated with phosphorylated STAT3 in cultured DD fibroblasts***. [64] | 6 |
| *TF** | Up | 0.000025 | 0.010 | 3 | Serotransferrin; Responsible for the transport of iron from sites of absorption and heme degradation; iron levels are increased in the lung tissues of patients with idiopathic pulmonary fibrosis; exogenous iron increases human lung fibroblast proliferation and cytokine responses. *Local tissue iron deposition occurs in the early cellular stage of DD*. [125, 126] | 4, 7 |
| *USP8** | Down | 0.001398 | 0.074 | 1 | Ubiquitin carboxyl-terminal hydrolase 8; USP8 pathway is essential for WNT/β-catenin signaling. WNT signaling pathways are central in DD biology. ***USP8 is under-expressed in DD Fibroblasts by Genome-wide exon expression profiles***. USP8 may exert protective influences against aging. *DD prevalence is age-related*. [10, 58] | 2, 3 |
| *YWHAZ* | Down | 0.007679 | 0.191 | 4 | 14-3-3 protein zeta/delta; Increased expression in hypertrophic scars; hypertrophic scar shares some ECM, microvascular, and mechanical load effects with DD. *Pathway analysis associates YWHAZ with BRAF. BRAF inhibitors provoke DD-like clinical changes.* [109, 129, 142, 143] | 3 |

| **S5 Table. Functionally enriched pathways in the hypothesis-based analysis.** | | | | | |
| --- | --- | --- | --- | --- | --- |
| **Category** | **ID** | **Description** | **STR** | **FDR** | **Genes of matching proteins in this network** |
| GO Process | GO:0050896 | Response to stimulus | 0.38 | 0.0004 | *PLAT, COL1A1, CSNK1G2, STAT3, HPX, MAP2K1, SMAD1, CASP3, AOC3, DAB2, STAT1, DDR2, LCN2, POSTN, YWHAZ, USP8, TF, PRKCA, SFRP4, SERPINH1, CSMD1, KNG1* |
| GO Process | GO:0042221 | Response to chemical | 0.56 | 0.0004 | *PLAT, COL1A1, STAT3, HPX, MAP2K1, SMAD1, CASP3, AOC3, DAB2, STAT1, DDR2, POSTN, USP8, TF, PRKCA, SFRP4, SERPINH1* |
| GO Process | GO:0009725 | Response to hormone | 1.01 | 0.0005 | *PLAT, COL1A1, STAT3, CASP3, DAB2, STAT1, DDR2, USP8, SFRP4* |
| GO Process | GO:0010033 | Response to organic substance | 0.65 | 0.0008 | *PLAT, COL1A1, STAT3, HPX, SMAD1, CASP3, DAB2, STAT1, DDR2, POSTN, USP8, PRKCA, SFRP4, SERPINH1* |
| GO Process | GO:0045597 | Positive regulation of cell differentiation | 0.94 | 0.0010 | *COL1A1, STAT3, MAP2K1, SMAD1, DAB2, STAT1, DDR2, PRKCA, SFRP4* |
| GO Process | GO:0007166 | Cell surface receptor signaling pathway | 0.7 | 0.0016 | *PLAT, COL1A1, CSNK1G2, STAT3, HPX, SMAD1, CASP3, DAB2, STAT1, DDR2, TF, SFRP4* |
| GO Process | GO:0009719 | Response to endogenous stimulus | 0.8 | 0.0016 | *PLAT, COL1A1, STAT3, SMAD1, CASP3, DAB2, STAT1, DDR2, USP8, SFRP4* |
| GO Process | GO:0009888 | Tissue development | 0.74 | 0.0016 | *COL1A1, MAP2K1, SMAD1, CASP3, DAB2, STAT1, DDR2, POSTN, ACAN, SERPINH1, CSMD1* |
| GO Process | GO:0030177 | Positive regulation of WNT signaling pathway | 1.48 | 0.0016 | *COL1A1, CSNK1G2, DAB2, USP8, SFRP4* |
| GO Process | GO:0030199 | Collagen fibril organization | 1.76 | 0.0016 | *COL1A1, DDR2, ACAN, SERPINH1* |
| GO Process | GO:0065008 | Regulation of biological quality | 0.55 | 0.0016 | *PLAT, STAT3, HPX, SMAD1, CASP3, DAB2, STAT1, LCN2, YWHAZ, USP8, TF, PRKCA, SFRP4, CSMD1, KNG1* |
| GO Process | GO:0071363 | Cellular response to growth factor stimulus | 1.1 | 0.0016 | *COL1A1, STAT3, SMAD1, CASP3, DAB2, DDR2, USP8* |
| GO Process | GO:0071560 | Cellular response to transforming growth factor beta stimulus | 1.45 | 0.0016 | *COL1A1, STAT3, SMAD1, DAB2, DDR2* |
| GO Process | GO:0007154 | Cell communication | 0.45 | 0.0022 | *PLAT, COL1A1, CSNK1G2, STAT3, HPX, MAP2K1, SMAD1, CASP3, DAB2, STAT1, DDR2, POSTN, YWHAZ, USP8, TF, PRKCA, SFRP4* |
| GO Process | GO:0051239 | Regulation of multicellular organismal process | 0.61 | 0.0022 | *PLAT, COL1A1, STAT3, MAP2K1, SMAD1, DAB2, STAT1, DDR2, LCN2, TF, PRKCA, SFRP4, KNG1* |
| GO Process | GO:0071495 | Cellular response to endogenous stimulus | 0.84 | 0.0022 | *PLAT, COL1A1, STAT3, SMAD1, CASP3, DAB2, STAT1, DDR2, USP8* |
| GO Process | GO:0006950 | Response to stress | 0.55 | 0.0025 | *PLAT, COL1A1, STAT3, HPX, MAP2K1, SMAD1, CASP3, AOC3, STAT1, DDR2, LCN2, TF, SERPINH1, KNG1* |
| GO Process | GO:0009967 | Positive regulation of signal transduction | 0.75 | 0.0025 | *COL1A1, CSNK1G2, STAT3, HPX, MAP2K1, DAB2, DDR2, USP8, PRKCA, SFRP4* |
| GO Process | GO:0010646 | Regulation of cell communication | 0.55 | 0.0025 | *PLAT, COL1A1, CSNK1G2, STAT3, HPX, MAP2K1, DAB2, STAT1, DDR2, POSTN, YWHAZ, USP8, PRKCA, SFRP4* |
| GO Process | GO:0014070 | Response to organic cyclic compound | 0.9 | 0.0025 | *PLAT, COL1A1, STAT3, SMAD1, CASP3, DAB2, STAT1, USP8* |
| GO Process | GO:0023051 | Regulation of signaling | 0.55 | 0.0025 | *PLAT, COL1A1, CSNK1G2, STAT3, HPX, MAP2K1, DAB2, STAT1, DDR2, POSTN, YWHAZ, USP8, PRKCA, SFRP4* |
| GO Process | GO:0048583 | Regulation of response to stimulus | 0.51 | 0.0025 | *PLAT, COL1A1, CSNK1G2, STAT3, HPX, MAP2K1, DAB2, STAT1, DDR2, POSTN, YWHAZ, USP8, PRKCA, SFRP4, KNG1* |
| GO Process | GO:0007165 | Signal transduction | 0.46 | 0.0030 | *PLAT, COL1A1, CSNK1G2, STAT3, HPX, MAP2K1, SMAD1, CASP3, DAB2, STAT1, DDR2, YWHAZ, USP8, TF, PRKCA, SFRP4* |
| GO Process | GO:0060348 | Bone development | 1.3 | 0.0030 | *COL1A1, SMAD1, DDR2, SFRP4, SERPINH1* |
| GO Process | GO:0009966 | Regulation of signal transduction | 0.57 | 0.0032 | *COL1A1, CSNK1G2, STAT3, HPX, MAP2K1, DAB2, STAT1, DDR2, POSTN, YWHAZ, USP8, PRKCA, SFRP4* |
| GO Process | GO:0007167 | Enzyme-linked receptor protein signaling pathway | 0.97 | 0.0035 | *PLAT, COL1A1, STAT3, SMAD1, CASP3, DDR2, TF* |
| GO Process | GO:0090263 | Positive regulation of canonical Wnt signaling pathway | 1.51 | 0.0039 | *COL1A1, CSNK1G2, USP8, SFRP4* |
| GO Process | GO:0016310 | Phosphorylation | 0.85 | 0.0045 | *CSNK1G2, STAT3, HPX, MAP2K1, SMAD1, DDR2, YWHAZ, PRKCA* |
| GO Process | GO:0048584 | Positive regulation of response to stimulus | 0.65 | 0.0045 | *PLAT, COL1A1, CSNK1G2, STAT3, HPX, MAP2K1, DAB2, DDR2, USP8, PRKCA, SFRP4* |
| GO Process | GO:0060828 | Regulation of canonical Wnt signaling pathway | 1.23 | 0.0045 | *COL1A1, CSNK1G2, DAB2, USP8, SFRP4* |
| GO Process | GO:0070887 | Cellular response to chemical stimulus | 0.6 | 0.0045 | *PLAT, COL1A1, STAT3, HPX, SMAD1, CASP3, DAB2, STAT1, DDR2, POSTN, USP8, TF* |
| GO Process | GO:0051171 | Regulation of nitrogen compound metabolic process | 0.4 | 0.0049 | *PLAT, COL1A1, STAT3, HPX, MAP2K1, SMAD1, CASP3, DAB2, STAT1, DDR2, YWHAZ, USP8, TF, PRKCA, SFRP4, SERPINH1, KNG1* |
| GO Process | GO:0008285 | Negative regulation of cell population proliferation | 0.92 | 0.0051 | *STAT3, MAP2K1, SMAD1, CASP3, DAB2, STAT1, SFRP4* |
| GO Process | GO:0006468 | Protein phosphorylation | 0.91 | 0.0057 | *CSNK1G2, HPX, MAP2K1, SMAD1, DDR2, YWHAZ, PRKCA* |
| GO Process | GO:0009653 | Anatomical structure morphogenesis | 0.63 | 0.0057 | *COL1A1, STAT3, MAP2K1, SMAD1, CASP3, YWHAZ, ACAN, PRKCA, SFRP4, SERPINH1, CSMD1* |
| GO Process | GO:0030198 | Extracellular matrix organization | 1.19 | 0.0057 | *COL1A1, DDR2, POSTN, ACAN, SERPINH1* |
| GO Process | GO:0042592 | Homeostatic process | 0.74 | 0.0057 | *STAT3, HPX, SMAD1, CASP3, STAT1, LCN2, TF, SFRP4, CSMD1* |
| GO Process | GO:0048513 | Animal organ development | 0.54 | 0.0057 | *COL1A1, STAT3, MAP2K1, SMAD1, CASP3, STAT1, DDR2, YWHAZ, TF, ACAN, SFRP4, SERPINH1, CSMD1* |
| GO Process | GO:0048545 | Response to steroid hormone | 1.18 | 0.0057 | *PLAT, COL1A1, CASP3, DAB2, USP8* |
| GO Process | GO:0048731 | System development | 0.49 | 0.0057 | *COL1A1, STAT3, MAP2K1, SMAD1, CASP3, STAT1, DDR2, YWHAZ, TF, ACAN, PRKCA, SFRP4, SERPINH1, CSMD1* |
| GO Process | GO:0048856 | Anatomical structure development | 0.43 | 0.0057 | *COL1A1, STAT3, MAP2K1, SMAD1, CASP3, DAB2, STAT1, DDR2, POSTN, YWHAZ, TF, ACAN, PRKCA, SFRP4, SERPINH1, CSMD1* |
| GO Process | GO:0080090 | Regulation of primary metabolic process | 0.39 | 0.0057 | *PLAT, COL1A1, STAT3, HPX, MAP2K1, SMAD1, CASP3, DAB2, STAT1, DDR2, YWHAZ, USP8, TF, PRKCA, SFRP4, SERPINH1, KNG1* |
| GO Process | GO:0071407 | Cellular response to organic cyclic compound | 1.01 | 0.0065 | *PLAT, COL1A1, SMAD1, CASP3, STAT1, USP8* |
| GO Process | GO:0001501 | Skeletal system development | 1 | 0.0067 | *COL1A1, SMAD1, DDR2, ACAN, SFRP4, SERPINH1* |
| GO Process | GO:0019222 | Regulation of metabolic process | 0.36 | 0.0067 | *PLAT, COL1A1, STAT3, HPX, MAP2K1, SMAD1, CASP3, DAB2, STAT1, DDR2, LCN2, YWHAZ, USP8, TF, PRKCA, SFRP4, SERPINH1, KNG1* |
| GO Process | GO:0070372 | Regulation of ERK1 and ERK2 cascade | 1.16 | 0.0067 | *MAP2K1, DAB2, DDR2, YWHAZ, PRKCA* |
| GO Process | GO:0009605 | Response to external stimulus | 0.6 | 0.0070 | *COL1A1, HPX, MAP2K1, CASP3, STAT1, DDR2, LCN2, POSTN, TF, CSMD1, KNG1* |
| GO Process | GO:0051240 | Positive regulation of multicellular organismal process | 0.71 | 0.0077 | *PLAT, COL1A1, STAT3, MAP2K1, SMAD1, DAB2, STAT1, LCN2, PRKCA* |
| GO Process | GO:0007259 | Receptor signaling pathway via JAK-STAT | 1.7 | 0.0091 | *STAT3, HPX, STAT1* |
| GO Process | GO:2000147 | Positive regulation of cell motility | 0.97 | 0.0092 | *COL1A1, STAT3, DAB2, DDR2, TF, PRKCA* |
| GO Process | GO:0006826 | Iron ion transport | 1.69 | 0.0095 | *HPX, LCN2, TF* |
| GO Process | GO:0048518 | Positive regulation of biological process | 0.37 | 0.0095 | *PLAT, COL1A1, CSNK1G2, STAT3, HPX, MAP2K1, SMAD1, CASP3, DAB2, STAT1, DDR2, LCN2, USP8, TF, PRKCA, SFRP4, KNG1* |
| GO Process | GO:1903034 | Regulation of response to wounding | 1.33 | 0.0095 | *PLAT, MAP2K1, DDR2, KNG1* |
| GO Process | GO:0031960 | Response to corticosteroid | 1.31 | 0.0098 | *PLAT, COL1A1, CASP3, USP8* |
| GO Process | GO:0048260 | Positive regulation of receptor-mediated endocytosis | 1.67 | 0.0098 | *DAB2, TF, SFRP4* |
| GO Process | GO:0051216 | Cartilage development | 1.31 | 0.0098 | *COL1A1, SMAD1, ACAN, SERPINH1* |
| GO Process | GO:0060255 | Regulation of macromolecule metabolic process | 0.37 | 0.0098 | *PLAT, COL1A1, STAT3, HPX, MAP2K1, SMAD1, CASP3, DAB2, STAT1, DDR2, YWHAZ, USP8, TF, PRKCA, SFRP4, SERPINH1, KNG1* |
| GO Process | GO:0070371 | ERK1 and ERK2 cascade | 1.67 | 0.0098 | *MAP2K1, YWHAZ, TF* |
| GO Process | GO:0071310 | Cellular response to organic substance | 0.63 | 0.0098 | *PLAT, COL1A1, STAT3, HPX, SMAD1, CASP3, DAB2, STAT1, DDR2, USP8* |
| GO Process | GO:0071417 | Cellular response to organonitrogen compound | 0.95 | 0.0098 | *PLAT, COL1A1, STAT3, CASP3, STAT1, DDR2* |
| GO Process | GO:0035295 | Tube development | 0.83 | 0.0100 | *MAP2K1, SMAD1, CASP3, STAT1, YWHAZ, PRKCA, CSMD1* |
| GO Process | GO:0043434 | Response to peptide hormone | 1.08 | 0.0106 | *PLAT, COL1A1, STAT3, STAT1, DDR2* |
| GO Process | GO:0051716 | Cellular response to stimulus | 0.36 | 0.0107 | *PLAT, COL1A1, CSNK1G2, STAT3, HPX, MAP2K1, SMAD1, CASP3, DAB2, STAT1, DDR2, POSTN, YWHAZ, USP8, TF, PRKCA, SFRP4* |
| GO Process | GO:0072359 | Circulatory system development | 0.82 | 0.0112 | *COL1A1, MAP2K1, SMAD1, CASP3, YWHAZ, ACAN, PRKCA* |
| GO Process | GO:0048878 | Chemical homeostasis | 0.82 | 0.0113 | *STAT3, HPX, STAT1, LCN2, TF, SFRP4, CSMD1* |
| GO Process | GO:0006879 | Cellular iron ion homeostasis | 1.61 | 0.0118 | *HPX, LCN2, TF* |
| GO Process | GO:0042127 | Regulation of cell population proliferation | 0.66 | 0.0120 | *STAT3, MAP2K1, SMAD1, CASP3, DAB2, STAT1, DDR2, PRKCA, SFRP4* |
| GO Process | GO:0032964 | Collagen biosynthetic process | 2.23 | 0.0161 | *COL1A1, SERPINH1* |
| GO Process | GO:0034103 | Regulation of tissue remodeling | 1.53 | 0.0186 | *DDR2, TF, PRKCA* |
| GO Process | GO:0009887 | Animal organ morphogenesis | 0.78 | 0.0191 | *COL1A1, STAT3, MAP2K1, ACAN, SFRP4, SERPINH1, CSMD1* |
| GO Process | GO:0000165 | MAPK cascade | 1.19 | 0.0214 | *MAP2K1, SMAD1, YWHAZ, TF* |
| GO Process | GO:0006952 | Defense response | 0.69 | 0.0214 | *STAT3, HPX, SMAD1, AOC3, STAT1, LCN2, TF, KNG1* |
| GO Process | GO:0007169 | Transmembrane receptor protein tyrosine kinase signaling pathway | 1 | 0.0214 | *PLAT, COL1A1, STAT3, CASP3, DDR2* |
| GO Process | GO:0038063 | Collagen-activated tyrosine kinase receptor signaling pathway | 2.15 | 0.0214 | *COL1A1, DDR2* |
| GO Process | GO:0060333 | Interferon-gamma-mediated signaling pathway | 2.15 | 0.0214 | *HPX, STAT1* |
| GO Process | GO:0048705 | Skeletal system morphogenesis | 1.17 | 0.0238 | *COL1A1, ACAN, SFRP4, SERPINH1* |
| GO Process | GO:1901701 | Cellular response to oxygen-containing compound | 0.75 | 0.0246 | *PLAT, COL1A1, STAT3, STAT1, DDR2, POSTN, USP8* |
| GO Process | GO:1901678 | Iron coordination entity transport | 2.09 | 0.0253 | *HPX, LCN2* |
| GO Process | GO:0042981 | Regulation of apoptotic process | 0.67 | 0.0269 | *CASP3, DAB2, STAT1, DDR2, YWHAZ, PRKCA, SFRP4, KNG1* |
| GO Process | GO:0071375 | Cellular response to peptide hormone stimulus | 1.15 | 0.0269 | *PLAT, STAT3, STAT1, DDR2* |
| GO Process | GO:0051591 | Response to cAMP | 1.45 | 0.0275 | *PLAT, COL1A1, STAT1* |
| GO Process | GO:0030154 | Cell differentiation | 0.47 | 0.0303 | *COL1A1, STAT3, MAP2K1, SMAD1, CASP3, DAB2, STAT1, YWHAZ, TF, ACAN, SFRP4, SERPINH1* |
| GO Process | GO:0032870 | Cellular response to hormone stimulus | 0.94 | 0.0335 | *PLAT, STAT3, STAT1, DDR2, USP8* |
| GO Process | GO:0045639 | Positive regulation of myeloid cell differentiation | 1.41 | 0.0336 | *STAT3, STAT1, PRKCA* |
| GO Process | GO:0060349 | Bone morphogenesis | 1.41 | 0.0336 | *COL1A1, SFRP4, SERPINH1* |
| GO Process | GO:0044092 | Negative regulation of molecular function | 0.72 | 0.0347 | *CASP3, AOC3, DAB2, DDR2, SFRP4, SERPINH1, KNG1* |
| GO Process | GO:0042542 | Response to hydrogen peroxide | 1.4 | 0.0348 | *COL1A1, CASP3, STAT1* |
| GO Process | GO:0050793 | Regulation of developmental process | 0.54 | 0.0348 | *COL1A1, STAT3, MAP2K1, SMAD1, DAB2, STAT1, DDR2, YWHAZ, PRKCA, SFRP4* |
| GO Process | GO:1901655 | Cellular response to ketone | 1.4 | 0.0348 | *PLAT, POSTN, USP8* |
| GO Process | GO:1901700 | Response to oxygen-containing compound | 0.65 | 0.0348 | *PLAT, COL1A1, STAT3, CASP3, STAT1, DDR2, POSTN, USP8* |
| GO Process | GO:2000641 | Regulation of early endosome to late endosome transport | 1.98 | 0.0348 | *MAP2K1, DAB2* |
| GO Process | GO:0034097 | Response to cytokine | 0.81 | 0.0355 | *COL1A1, STAT3, HPX, CASP3, STAT1, PRKCA* |
| GO Process | GO:0048522 | Positive regulation of cellular process | 0.36 | 0.0355 | *COL1A1, CSNK1G2, STAT3, HPX, MAP2K1, SMAD1, CASP3, DAB2, STAT1, DDR2, USP8, TF, PRKCA, SFRP4, KNG1* |
| GO Process | GO:0045780 | Positive regulation of bone resorption | 1.95 | 0.0361 | *TF, PRKCA* |
| GO Process | GO:0033993 | Response to lipid | 0.79 | 0.0397 | *PLAT, COL1A1, STAT3, CASP3, DAB2, USP8* |
| GO Process | GO:0030335 | Positive regulation of cell migration | 0.91 | 0.0416 | *COL1A1, STAT3, DAB2, DDR2, PRKCA* |
| GO Process | GO:0045765 | Regulation of angiogenesis | 1.08 | 0.0429 | *STAT3, SMAD1, STAT1, PRKCA* |
| GO Process | GO:0065007 | Biological regulation | 0.18 | 0.0433 | *PLAT, COL1A1, CSNK1G2, STAT3, HPX, MAP2K1, SMAD1, CASP3, AOC3, DAB2, STAT1, DDR2, LCN2, POSTN, YWHAZ, USP8, TF, PRKCA, SFRP4, SERPINH1, CSMD1, KNG1* |
| GO Process | GO:0071295 | Cellular response to vitamin | 1.89 | 0.0445 | *COL1A1, POSTN* |
| GO Process | GO:0032501 | Multicellular organismal process | 0.32 | 0.0448 | *PLAT, COL1A1, STAT3, MAP2K1, SMAD1, CASP3, STAT1, DDR2, YWHAZ, TF, ACAN, PRKCA, SFRP4, SERPINH1, CSMD1, KNG1* |
| GO Process | GO:0051246 | Regulation of protein metabolic process | 0.51 | 0.0472 | *PLAT, STAT3, HPX, MAP2K1, CASP3, DAB2, DDR2, USP8, SERPINH1, KNG1* |
| GO Process | GO:0032355 | Response to estradiol | 1.33 | 0.0476 | *COL1A1, STAT3, CASP3* |
| GO Process | GO:0042325 | Regulation of phosphorylation | 0.68 | 0.0498 | *STAT3, HPX, MAP2K1, CASP3, DAB2, DDR2, TF* |
| GO Process | GO:2000637 | Positive regulation of miRNA-mediated gene silencing | 1.85 | 0.0498 | *STAT3, MAP2K1* |
| GO Component | GO:0062023 | Collagen-containing extracellular matrix | 1.1 | 0.0124 | *COL1A1, HPX, POSTN, ACAN, SERPINH1, KNG1* |
| GO Component | GO:0072562 | Blood microparticle | 1.46 | 0.0124 | *HPX, YWHAZ, TF, KNG1* |
| GO Component | GO:0031410 | Cytoplasmic vesicle | 0.58 | 0.0228 | *PLAT, COL1A1, HPX, MAP2K1, AOC3, DAB2, LCN2, YWHAZ, USP8, TF, KNG1* |
| GO Component | GO:0071944 | Cell periphery | 0.36 | 0.0391 | *COL1A1, CSNK1G2, STAT3, HPX, MAP2K1, CASP3, AOC3, DAB2, DDR2, POSTN, USP8, TF, ACAN, PRKCA, SERPINH1, KNG1* |
| GO Component | GO:0012505 | Endomembrane system | 0.4 | 0.0453 | *PLAT, COL1A1, MAP2K1, SMAD1, AOC3, DAB2, LCN2, POSTN, USP8, TF, ACAN, PRKCA, SERPINH1, KNG1* |
| KEGG | hsa05161 | Hepatitis B | 1.51 | 9.10E-06 | *STAT3, MAP2K1, CASP3, STAT1, YWHAZ, PRKCA* |
| KEGG | hsa04933 | AGE-RAGE signaling pathway in diabetic complications | 1.65 | 1.67E-05 | *COL1A1, STAT3, CASP3, STAT1, PRKCA* |
| KEGG | hsa05160 | Hepatitis C | 1.44 | 0.0001 | *STAT3, MAP2K1, CASP3, STAT1, YWHAZ* |
| KEGG | hsa05205 | Proteoglycans in cancer | 1.34 | 0.0002 | *COL1A1, STAT3, MAP2K1, CASP3, PRKCA* |
| KEGG | hsa04066 | HIF-1 signaling pathway | 1.53 | 0.0004 | *STAT3, MAP2K1, TF, PRKCA* |
| KEGG | hsa04935 | Growth hormone synthesis, secretion and action | 1.47 | 0.0006 | *STAT3, MAP2K1, STAT1, PRKCA* |
| KEGG | hsa05200 | Pathways in cancer | 1 | 0.0011 | *STAT3, MAP2K1, CASP3, STAT1, PRKCA, KNG1* |
| KEGG | hsa05164 | Influenza A | 1.32 | 0.0015 | *MAP2K1, CASP3, STAT1, PRKCA* |
| KEGG | hsa05206 | MicroRNAs in cancer | 1.33 | 0.0015 | *STAT3, MAP2K1, CASP3, PRKCA* |
| KEGG | hsa05167 | Kaposi sarcoma-associated herpesvirus infection | 1.26 | 0.0022 | *STAT3, MAP2K1, CASP3, STAT1* |
| KEGG | hsa04917 | Prolactin signaling pathway | 1.58 | 0.0023 | *STAT3, MAP2K1, STAT1* |
| KEGG | hsa05212 | Pancreatic cancer | 1.56 | 0.0023 | *STAT3, MAP2K1, STAT1* |
| KEGG | hsa05223 | Non-small cell lung cancer | 1.58 | 0.0023 | *STAT3, MAP2K1, PRKCA* |
| KEGG | hsa01521 | EGFR tyrosine kinase inhibitor resistance | 1.52 | 0.0026 | *STAT3, MAP2K1, PRKCA* |
| KEGG | hsa05163 | Human cytomegalovirus infection | 1.2 | 0.0026 | *STAT3, MAP2K1, CASP3, PRKCA* |
| KEGG | hsa05235 | PD-L1 expression and PD-1 checkpoint pathway in cancer | 1.47 | 0.0032 | *STAT3, MAP2K1, STAT1* |
| KEGG | hsa05145 | Toxoplasmosis | 1.4 | 0.0046 | *STAT3, CASP3, STAT1* |
| KEGG | hsa05146 | Amoebiasis | 1.41 | 0.0046 | *COL1A1, CASP3, PRKCA* |
| KEGG | hsa04726 | Serotonergic synapse | 1.38 | 0.0050 | *MAP2K1, CASP3, PRKCA* |
| KEGG | hsa04071 | Sphingolipid signaling pathway | 1.35 | 0.0058 | *MAP2K1, PRKCA, KNG1* |
| KEGG | hsa04650 | Natural killer cell mediated cytotoxicity | 1.33 | 0.0061 | *MAP2K1, CASP3, PRKCA* |
| KEGG | hsa04919 | Thyroid hormone signaling pathway | 1.33 | 0.0061 | *MAP2K1, STAT1, PRKCA* |
| KEGG | hsa04926 | Relaxin signaling pathway | 1.31 | 0.0064 | *COL1A1, MAP2K1, PRKCA* |
| KEGG | hsa05165 | Human papillomavirus infection | 1.02 | 0.0073 | *COL1A1, MAP2K1, CASP3, STAT1* |
| KEGG | hsa05162 | Measles | 1.27 | 0.0075 | *STAT3, CASP3, STAT1* |
| KEGG | hsa04550 | Signaling pathways regulating pluripotency of stem cells | 1.26 | 0.0078 | *STAT3, MAP2K1, SMAD1* |
| KEGG | hsa04151 | PI3K-Akt signaling pathway | 0.99 | 0.0085 | *COL1A1, MAP2K1, YWHAZ, PRKCA* |
| KEGG | hsa05143 | African trypanosomiasis | 1.68 | 0.0107 | *PRKCA, KNG1* |
| KEGG | hsa05203 | Viral carcinogenesis | 1.15 | 0.0147 | *STAT3, CASP3, YWHAZ* |
| KEGG | hsa04062 | Chemokine signaling pathway | 1.14 | 0.0149 | *STAT3, MAP2K1, STAT1* |
| KEGG | hsa05169 | Epstein-Barr virus infection | 1.13 | 0.0158 | *STAT3, CASP3, STAT1* |
| KEGG | hsa04510 | Focal adhesion | 1.12 | 0.0160 | *COL1A1, MAP2K1, PRKCA* |
| KEGG | hsa05170 | Human immunodeficiency virus 1 infection | 1.1 | 0.0174 | *MAP2K1, CASP3, PRKCA* |
| KEGG | hsa04370 | VEGF signaling pathway | 1.49 | 0.0204 | *MAP2K1, PRKCA* |
| KEGG | hsa04730 | Long-term depression | 1.46 | 0.0219 | *MAP2K1, PRKCA* |
| KEGG | hsa05321 | Inflammatory bowel disease | 1.46 | 0.0219 | *STAT3, STAT1* |
| KEGG | hsa04720 | Long-term potentiation | 1.43 | 0.0235 | *MAP2K1, PRKCA* |
| KEGG | hsa04929 | GnRH secretion | 1.43 | 0.0235 | *MAP2K1, PRKCA* |
| KEGG | hsa04664 | Fc epsilon RI signaling pathway | 1.42 | 0.0237 | *MAP2K1, PRKCA* |
| KEGG | hsa05221 | Acute myeloid leukemia | 1.41 | 0.0244 | *STAT3, MAP2K1* |
| KEGG | hsa05214 | Glioma | 1.38 | 0.0266 | *MAP2K1, PRKCA* |
| KEGG | hsa04010 | MAPK signaling pathway | 0.95 | 0.0334 | *MAP2K1, CASP3, PRKCA* |
| KEGG | hsa04012 | ErbB signaling pathway | 1.33 | 0.0334 | *MAP2K1, PRKCA* |
| KEGG | hsa04610 | Complement and coagulation cascades | 1.32 | 0.0334 | *PLAT, KNG1* |
| KEGG | hsa05210 | Colorectal cancer | 1.32 | 0.0334 | *MAP2K1, CASP3* |
| KEGG | hsa04540 | Gap junction | 1.29 | 0.0350 | *MAP2K1, PRKCA* |
| KEGG | hsa04912 | GnRH signaling pathway | 1.29 | 0.0350 | *MAP2K1, PRKCA* |
| KEGG | hsa04657 | IL-17 signaling pathway | 1.27 | 0.0358 | *CASP3, LCN2* |
| KEGG | hsa04666 | Fc gamma R-mediated phagocytosis | 1.28 | 0.0358 | *MAP2K1, PRKCA* |
| KEGG | hsa04750 | Inflammatory mediator regulation of TRP channels | 1.27 | 0.0358 | *PRKCA, KNG1* |
| KEGG | hsa04916 | Melanogenesis | 1.26 | 0.0373 | *MAP2K1, PRKCA* |
| KEGG | hsa05231 | Choline metabolism in cancer | 1.26 | 0.0373 | *MAP2K1, PRKCA* |
| KEGG | hsa05215 | Prostate cancer | 1.25 | 0.0374 | *PLAT, MAP2K1* |
| KEGG | hsa04620 | Toll-like receptor signaling pathway | 1.23 | 0.0381 | *MAP2K1, STAT1* |
| KEGG | hsa04659 | Th17 cell differentiation | 1.24 | 0.0381 | *STAT3, STAT1* |
| KEGG | hsa04928 | Parathyroid hormone synthesis, secretion and action | 1.22 | 0.0404 | *MAP2K1, PRKCA* |
| KEGG | hsa04725 | Cholinergic synapse | 1.2 | 0.0433 | *MAP2K1, PRKCA* |
| KEGG | hsa04668 | TNF signaling pathway | 1.19 | 0.0441 | *MAP2K1, CASP3* |
| Reactome | HSA-1474244 | Extracellular matrix organization | 1.23 | 0.0025 | *COL1A1, CASP3, DDR2, ACAN, PRKCA, SERPINH1* |
| Reactome | HSA-9006934 | Signaling by Receptor Tyrosine Kinases | 1.06 | 0.0025 | *PLAT, COL1A1, STAT3, MAP2K1, STAT1, USP8, PRKCA* |
| Reactome | HSA-162582 | Signal Transduction | 0.61 | 0.0055 | *PLAT, COL1A1, CSNK1G2, STAT3, MAP2K1, SMAD1, CASP3, STAT1, YWHAZ, USP8, PRKCA, KNG1* |
| Reactome | HSA-449147 | Signaling by Interleukins | 1.05 | 0.0063 | *STAT3, MAP2K1, CASP3, STAT1, LCN2, YWHAZ* |
| Reactome | HSA-76002 | Platelet activation, signaling and aggregation | 1.22 | 0.0063 | *COL1A1, YWHAZ, TF, PRKCA, KNG1* |
| Reactome | HSA-1433557 | Signaling by SCF-KIT | 1.78 | 0.0077 | *STAT3, STAT1, PRKCA* |
| Reactome | HSA-109606 | Intrinsic Pathway for Apoptosis | 1.69 | 0.0114 | *STAT3, CASP3, YWHAZ* |
| Reactome | HSA-109582 | Hemostasis | 0.93 | 0.0136 | *PLAT, COL1A1, YWHAZ, TF, PRKCA, KNG1* |
| Reactome | HSA-186797 | Signaling by PDGF | 1.65 | 0.0136 | *PLAT, STAT3, STAT1* |
| Reactome | HSA-3000171 | Non-integrin membrane-ECM interactions | 1.64 | 0.0136 | *COL1A1, DDR2, PRKCA* |
| Reactome | HSA-8985947 | Interleukin-9 signaling | 2.28 | 0.0148 | *STAT3, STAT1* |
| Reactome | HSA-9020958 | Interleukin-21 signaling | 2.23 | 0.0162 | *STAT3, STAT1* |
| Reactome | HSA-1059683 | Interleukin-6 signaling | 2.19 | 0.0177 | *STAT3, STAT1* |
| Reactome | HSA-430116 | GP1b-IX-V activation signaling | 2.15 | 0.0177 | *COL1A1, YWHAZ* |
| Reactome | HSA-6806834 | Signaling by MET | 1.51 | 0.0177 | *COL1A1, STAT3, USP8* |
| Reactome | HSA-8984722 | Interleukin-35 Signaling | 2.15 | 0.0177 | *STAT3, STAT1* |
| Reactome | HSA-9020956 | Interleukin-27 signaling | 2.19 | 0.0177 | *STAT3, STAT1* |
| Reactome | HSA-9673767 | Signaling by PDGFRA transmembrane, juxtamembrane and kinase domain mutants | 2.15 | 0.0177 | *STAT3, STAT1* |
| Reactome | HSA-9673770 | Signaling by PDGFRA extracellular domain mutants | 2.15 | 0.0177 | *STAT3, STAT1* |
| Reactome | HSA-1839117 | Signaling by cytosolic FGFR1 fusion mutants | 1.98 | 0.0265 | *STAT3, STAT1* |
| Reactome | HSA-6785807 | Interleukin-4 and Interleukin-13 signaling | 1.38 | 0.0286 | *STAT3, STAT1, LCN2* |
| Reactome | HSA-9670439 | Signaling by phosphorylated juxtamembrane, extracellular and kinase domain KIT mutants | 1.93 | 0.0294 | *STAT3, STAT1* |
| Reactome | HSA-195721 | Signaling by WNT | 1.06 | 0.0328 | *CSNK1G2, YWHAZ, USP8, PRKCA* |
| Reactome | HSA-8878166 | Transcriptional regulation by RUNX2 | 1.33 | 0.0328 | *COL1A1, SMAD1, STAT1* |
| Reactome | HSA-9705462 | Inactivation of CSF3 (G-CSF) signaling | 1.85 | 0.0328 | *STAT3, STAT1* |
| Reactome | HSA-982772 | Growth hormone receptor signaling | 1.87 | 0.0328 | *STAT3, STAT1* |
| Reactome | HSA-8854691 | Interleukin-20 family signaling | 1.84 | 0.0330 | *STAT3, STAT1* |
| Reactome | HSA-76005 | Response to elevated platelet cytosolic Ca2+ | 1.29 | 0.0351 | *TF, PRKCA, KNG1* |
| Reactome | HSA-3000170 | Syndecan interactions | 1.8 | 0.0358 | *COL1A1, PRKCA* |
| Reactome | HSA-1474228 | Degradation of the extracellular matrix | 1.26 | 0.0398 | *COL1A1, CASP3, ACAN* |
| Reactome | HSA-186763 | Downstream signal transduction | 1.77 | 0.0398 | *STAT3, STAT1* |
| Reactome | HSA-8941326 | RUNX2 regulates bone development | 1.74 | 0.0415 | *COL1A1, SMAD1* |
| Reactome | HSA-9680350 | Signaling by CSF1 (M-CSF) in myeloid cells | 1.74 | 0.0415 | *STAT3, STAT1* |

S5 Table. Functionally enriched pathways derived from the 23 differentially expressed proteins in the hypothesis-based analysis. ID: Enriched category name. STR: Strength of protein-protein interaction (PPI). FDR: False Discovery Rate (Adjusted p-value). Overall PPI enrichment p-value: 1.35E-07. Hypothesis-based candidate protein preselection may influence the enrichment results. Figure 6 summarizes enriched pathways for individual genes.

| **S6 Table. Genes referenced in the manuscript and their aliases** | | |
| --- | --- | --- |
| **Gene** | **UniProt** | **Proteins and gene aliases (Separated by ";" Character)** |
| *ACAN* | P16112 | *ACAN*; aggrecan; CSPGCP; aggrecan proteoglycan; chondroitin sulfate proteoglycan 1; aggrecan 1; AGC1; AGCAN; CSPG1; SEDK; SSOAOD; cartilage-specific proteoglycan core protein; chondroitin sulfate proteoglycan core protein 1; large aggregating proteoglycan; Aggrecan core protein; CSPCP |
| *AKR1A1* | P14550 | *AKR1A1*; aldo-keto reductase family 1 member A1; ALR; DD3; dihydrodiol dehydrogenase 3; aldo-keto reductase family 1, member A1 (aldehyde reductase); HEL-S-6; HEL-S-165mP; epididymis secretory protein Li 6; epididymis secretory sperm binding protein Li 165mP; glucuronate reductase; glucuronolactone reductase; Alcohol dehydrogenase [NADP(+)] |
| *AOC3* | Q16853 | *AOC3*; amine oxidase copper containing 3; VAP1; HPAO; VAP-1; vascular adhesion protein 1; SSAO; membrane primary amine oxidase; amine oxidase, copper containing 3 (vascular adhesion protein 1); copper amine oxidase; placenta copper monamine oxidase; semicarbazide-sensitive amine oxidase |
| *ARG2* | P78540 | *ARG2*; Arginase 2; Arginase-2 Mitochondrial; Kidney-Type Arginase; Non-Hepatic Arginase; Arginase Type II; Type II Arginase; Arginase II; EC 3.5.3.1; L-Arginine Amidinohydrolase; L-Arginine Ureahydrolase; Nonhepatic Arginase; Kidney Arginase; EC 3.5.3 |
| *BIN2* | Q9UBW5 | *BIN2*; Bridging Integrator 2; BRAP-1; Breast Cancer-Associated Protein 1; Breast Cancer Associated Protein BRAP1; BRAP1 |
| *C5* | P01031 | *C5*; complement C5; CPAMD4; C5a; C5b; prepro-C5; C5a anaphylatoxin; complement component 5; C5D; ECLZB; C3 and PZP-like alpha-2-macroglobulin domain-containing protein 4; anaphylatoxin C5a analog |
| *C6* | P13671 | *C6*; complement C6; complement component 6; complement component C6 |
| *CALCB* | P10092 | *CALCB*; Calcitonin Related Polypeptide Beta; CGRP-II; CALC2; Calcitonin Gene-Related Peptide II; Calcitonin Gene-Related Peptide 2; Beta-Type CGRP; Calcitonin 2; Beta-CGRP; FLJ30166; CGRP2 |
| *CASP3* | P42574 | *CASP3*; caspase 3; CPP32; CPP32B; Yama; apopain; caspase 3, apoptosis-related cysteine protease; caspase 3, apoptosis-related cysteine peptidase; SCA-1; caspase-3; CASP-3; CPP-32; PARP cleavage protease; SREBP cleavage activity 1; cysteine protease CPP32; procaspase3; protein Yama |
| *CGA* | P01215 | *CGA*; glycoprotein hormones, alpha polypeptide; HCG; GPHa; GPHA1; FSHA; LHA; TSHA; GPA1; follicle-stimulating hormone alpha subunit; chorionic gonadotropin, alpha polypeptide; luteinizing hormone alpha chain; lutropin alpha chain; thyroid-stimulating hormone alpha chain; glycoprotein hormones alpha chain; CG-ALPHA; FSH-alpha; LSH-alpha; TSH-alpha; anterior pituitary glycoprotein hormones common subunit alpha; choriogonadotropin alpha chain; chorionic gonadotrophin subunit alpha; follicle-stimulating hormone alpha chain; follitropin alpha chain; thyrotropin alpha chain |
| *CGB3* | P0DN86 | *CGB3*; Chorionic Gonadotropin Subunit Beta 3; CGB; Chorionic Gonadotropin Beta Polypeptide; Chorionic Gonadotropin Beta Subunit 3; Choriogonadotropin Subunit Beta 3; Chorionic Gonadotropin Chain Beta; CG-Beta; Chorionic Gonadotropin Beta 3 Subunit; Chorionic Gonadotrophin Chain Beta; Chorionic Gonadotropin Beta Chain; Luteinizing Hormone Beta Subunit; Choriogonadotropin Subunit Beta; CGB5; CGB7; CGB8; HCGB; LHB |
| *CGB7* | P0DN87 | *CGB7*; Chorionic Gonadotropin Subunit Beta 7; CG-Beta-A; Chorionic Gonadotropin Beta Polypeptide 7; Chorionic Gonadotropin Beta Subunit 7; Choriogonadotropin Subunit Beta 7; Chorionic Gonadotropin Beta 7 Subunit; CGB6 |
| *COL1A1* | P02452 | *COL1A1*; collagen type I alpha 1 chain; collagen, type I, alpha 1; CAFYD; EDSARTH1; OI1; OI2; OI3; collagen alpha-1(I) chain; alpha1(I) procollagen; collagen alpha 1 chain type I; collagen alpha-1(I) chain preproprotein; collagen of skin, tendon and bone, alpha-1 chain; pro-alpha-1 collagen type 1; type I proalpha 1; type I procollagen alpha 1 chain; Alpha-1 type I collagen |
| *CPB1* | P15086 | *CPB1*; Carboxypeptidase B1; Pancreatic Carboxypeptidase B; Carboxypeptidase B1 (Tissue); Tissue Carboxypeptidase B; Pancreas-Specific Protein; Carboxypeptidase B; Protaminase; EC 3.4.17.2; PASP; PCPB; CPB; Procarboxypeptidase B; EC 3.4.17 |
| *CRKL* | P46109 | *CRKL*; CRK Like Proto-Oncogene; V-Crk Avian Sarcoma Virus CT10 Oncogene Homolog-Like; Crk-Like Protein |
| *CSMD1* | Q96PZ7 | *CSMD1*; CUB and Sushi multiple domains 1; KIAA1890; PPP1R24; protein phosphatase 1, regulatory subunit 24; CUB and sushi domain-containing protein 1; CUB and sushi multiple domains protein 1 |
| *CSNK1G2* | P78368 | *CSNK1G2*; Casein Kinase 1 Gamma 2; CK1g2; Casein Kinase I Isoform Gamma-2; CKI-Gamma 2; Casein Kinase 1 Gamma 2; Casein Kinase I; CK1G2 |
| *DAB2* | P98082 | *DAB2*; DAB adaptor protein 2; DOC-2; disabled (Drosophila) homolog 2 (mitogen-responsive phosphoprotein); disabled homolog 2, mitogen-responsive phosphoprotein (Drosophila); Dab, mitogen-responsive phosphoprotein, homolog 2 (Drosophila); DAB2, clathrin adaptor protein; DOC2; disabled homolog 2; Dab, mitogen-responsive phosphoprotein, homolog 2; adaptor molecule disabled-2; differentially expressed in ovarian carcinoma 2; differentially-expressed protein 2; disabled homolog 2, mitogen-responsive phosphoprotein |
| *DDR2* | Q16832 | *DDR2*; discoidin domain receptor tyrosine kinase 2; TKT; discoidin domain receptor family, member 2; MIG20a; NTRKR3; TYRO10; WRCN; discoidin domain-containing receptor 2; CD167 antigen-like family member B; cell migration-inducing protein 20; discoidin domain receptor 2; discoidin domain-containing receptor tyrosine kinase 2; migration-inducing gene 16 protein; neurotrophic tyrosine kinase receptor related 3; receptor protein-tyrosine kinase TKT; tyrosine-protein kinase TYRO10; Neurotrophic tyrosine kinase, receptor-related 3; CD167b antigen |
| *DDX19A* | Q9NUU7 | *DDX19A*; DEAD-Box Helicase 19A; DDX19L; DEAD (Asp-Glu-Ala-Asp) Box Polypeptide 19A; ATP-Dependent RNA Helicase DDX19A; DEAD Box Protein 19A; DDX19-Like Protein; FLJ11126; DEAD (Asp-Glu-Ala-As) Box Polypeptide 19-Like; DEAD (Asp-Glu-Ala-As) Box Polypeptide 19A; DDX19-DDX19L; RNA Helicase; EC 3.6.4.13; EC 3.6.1 |
| *DSG2* | Q14126 | *DSG2*; Desmoglein 2; CDHF5; Cadherin Family Member 5; Desmoglein-2; HDGC |
| *EDIL3* | O43854 | *EDIL3*; EGF Like Repeats And Discoidin Domains 3; DEL1; EGF-Like Repeat And Discoidin I-Like Domain-Containing Protein 3; Developmentally-Regulated Endothelial Cell Locus 1 Protein; EGF-Like Repeats And Discoidin I-Like Domains 3; Integrin-Binding Protein DEL1; Developmental Endothelial Locus-1 |
| *EIF4H* | Q15056 | *EIF4H*; Eukaryotic Translation Initiation Factor 4H; WSCR1; KIAA0038; WBSCR1; Williams-Beuren Syndrome Chromosome Region 1; EIF-4H; Williams-Beuren Syndrome Chromosomal Region 1 Protein |
| *ELAPOR1* | Q6UXG2 | *ELAPOR1*; Endosome-Lysosome Associated Apoptosis And Autophagy Regulator 1; KIAA1324; EIG121; Endosome/Lysosome-Associated Apoptosis And Autophagy Regulator 1; Estrogen-Induced Gene 121 Protein; Maba1; Estrogen Induced Gene 121; MABA1 |
| *FAM234B* | A2RU67 | *FAM234B*; Family With Sequence Similarity 234 Member B; KIAA1467; Protein FAM234B |
| *FKBP5* | Q13451 | *FKBP5*; FKBP Prolyl Isomerase 5; FKBP51; FKBP54; P54; 54 KDa Progesterone Receptor-Associated Immunophilin; Peptidyl-Prolyl Cis-Trans Isomerase FKBP5; 51 KDa FK506-Binding Protein; Androgen-Regulated Protein 6; HSP90-Binding Immunophilin; FK506-Binding Protein 5; FK506 Binding Protein 5; PPIase FKBP5; 51 KDa FKBP; FF1 Antigen; EC 5.2.1.8; Rotamase; FKBP-51; PPIase; Ptg-10; AIG6; Peptidylprolyl Cis-Trans Isomerase; T-Cell FK506-Binding Protein; PPIASE; PTG-10; FKBP-5 |
| *G3BP2* | Q9UN86 | *G3BP2*; G3BP Stress Granule Assembly Factor 2; KIAA0660; Ras-GTPase Activating Protein SH3 Domain-Binding Protein 2; GTPase Activating Protein (SH3 Domain) Binding Protein 2; Ras GTPase-Activating Protein-Binding Protein 2; GAP SH3 Domain-Binding Protein 2; G3BP-2 |
| *GP1BB* | P13224 | *GP1BB*; Glycoprotein Ib Platelet Subunit Beta; Platelet Glycoprotein Ib Beta Chain; Glycoprotein Ib (Platelet) Beta Polypeptide; Antigen CD42b-Beta; GP-Ib Beta; GPIbbeta; CD42C; Nuclear Localization Signal Deleted In Velocardiofacial Syndrome; Platelet Membrane Glycoprotein Ib Beta; Glycoprotein Ib Platelet Beta Subunit; CD42c Antigen; GPIb-Beta; GPIBBETA; BDPLT1; CD42c; GPIBB; GPIbB; BS |
| *GUSB* | P08236 | *GUSB*; glucuronidase beta; glucuronidase, beta; BG; MPS7; beta-glucuronidase; beta-D-glucuronidase; beta-G1 |
| *HGS* | O14964 | *HGS*; Hepatocyte Growth Factor-Regulated Tyrosine Kinase Substrate; Protein Pp110; ZFYVE8; Human Growth Factor-Regulated Tyrosine Kinase Substrate; Vps27; VPS27 |
| *HPX* | P02790 | *HPX*; hemopexin; HX; beta-1B-glycoprotein |
| *HSP90AA1* | P07900 | *HSP90AA1*; Heat Shock Protein 90 Alpha Family Class A Member 1; HSP90N; HSPC1; HSPCA; Heat Shock Protein 90kDa Alpha (Cytosolic) Class A Member 1; Lipopolysaccharide-Associated Protein 2; Heat Shock 90kDa Protein 1 Alpha; Renal Carcinoma Antigen NY-REN-38; Heat Shock 90kD Protein 1 Alpha; Heat Shock Protein HSP 90-Alpha; LPS-Associated Protein 2; Heat Shock 86 KDa; FLJ31884; HSP90A; HSP 86; Hsp89; Hsp90; HSP86; LAP-2; Heat Shock Protein 90kDa Alpha Family Class A Member 1; Epididymis Secretory Sperm Binding Protein Li 65p; Epididymis Luminal Secretory Protein 52; Heat Shock 90kD Protein 1 Alpha-Like 4; Heat Shock 90kD Protein Alpha-Like 4; EC 3.6.4.10; HEL-S-65p; HSPCAL1; HSPCAL4; HSP89A; Hsp103; HSP89; HSP90; EL52; HSPN; LAP2 |
| *KNG1* | P01042 | *KNG1*; kininogen 1; BK; HMWK; alpha-2-thiol proteinase inhibitor; bradykinin; high-molecular-weight kininogen; kininogen; BDK; HAE6; HK; KNG; kininogen-1; Fitzgerald factor; high molecular weight kininogen; Williams-Fitzgerald-Flaujeac factor |
| *LCN2* | P80188 | *LCN2*; Lipocalin 2; NGAL; Neutrophil Gelatinase-Associated Lipocalin; Oncogene 24p3; 24p3; 25 KDa Alpha-2-Microglobulin-Related Subunit Of MMP-9; Siderocalin; P25; Migration-Stimulating Factor Inhibitor; Lipocalin 2 (Oncogene 24p3); Siderocalin LCN2; Lipocalin-2; MSFI; HNL |
| *MAP2K1* | Q02750 | *MAP2K1*; mitogen-activated protein kinase kinase 1; MEK1; MAPKK1; CFC3; MEL; MKK1; PRKMK1; dual specificity mitogen-activated protein kinase kinase 1; ERK activator kinase 1; MAPK/ERK kinase 1; MAPKK 1; MEK 1; protein kinase, mitogen-activated, kinase 1 (MAP kinase kinase 1); MAP kinase kinase 1 |
| *MAP3K11* | Q16584 | *MAP3K11*; Mitogen-Activated Protein Kinase Kinase Kinase 11; SPRK; MEKK11; MLK3; PTK1; Src-Homology 3 Domain-Containing Proline-Rich Kinase; Mixed Lineage Kinase 3; EC 2.7.11.25; SH3 Domain-Containing Proline-Rich Kinase; Protein-Tyrosine Kinase PTK1; EC 2.7.11; MLK-3 |
| *MBNL1* | Q9NR56 | *MBNL1*; Muscleblind-Like Protein 1; EXP42; EXP40; EXP35; Muscleblind-Like Splicing Regulator 1; Muscleblind (Drosophila)-Like; Muscleblind-Like (Drosophila); Muscleblind-Like |
| *MBNL2* | Q5VZF2 | *MBNL2*; Muscleblind Like Splicing Regulator 2; MBLL39; MBLL; Muscleblind-Like Protein-Like 39; Muscleblind-Like Protein 2; Muscleblind-Like Protein 1; Muscleblind-Like Splicing Regulator 2; Muscleblind-Like 2 (Drosophila); Muscleblind-Like Protein-Like; Muscleblind-Like 2; PRO2032; MLP1 |
| *NAA80* | Q93015 | *NAA80*; N-Alpha-Acetyltransferase 80; NatH Catalytic Subunit; FUS2; NAT6; N-Alpha-Acetyltransferase 80; N-Acetyltransferase 6; Protein Fusion-2; HsNAAA80; N(Alpha)-Acetyltransferase 80 NatH Catalytic Subunit; N-Acetyltransferase 6 (GCN5-Related); Protein Fus-2; EC 2.3.1.-; FUS-2 |
| *NRG4* | Q8WWG1 | *NRG4*; Neuregulin 4; HRG4; Pro-Neuregulin-4 Membrane-Bound Isoform; Pro-NRG4; Heregulin 4 |
| *OSCAR* | Q8IYS5 | *OSCAR*; Osteoclast Associated Ig-Like Receptor; Osteoclast Associated Immunoglobulin-Like Receptor; Osteoclast-Associated Immunoglobulin-Like Receptor; Polymeric Immunoglobulin Receptor 3; Poly-Ig Receptor 3; PIgR-3; Osteoclast Associated Receptor OSCAR-S1; Osteoclast Associated Receptor OSCAR-S2; Osteoclast-Associated Receptor; HOSCAR; PIGR3; PIgR3 |
| *PCSK9* | Q8NBP7 | *PCSK9*; proprotein convertase subtilisin/kexin type 9; NARC-1; FH3; hypercholesterolemia, autosomal dominant 3; FHCL3; HCHOLA3; LDLCQ1; NARC1; PC9; convertase subtilisin/kexin type 9 preproprotein; neural apoptosis regulated convertase 1; subtilisin/kexin-like protease PC9; Neural apoptosis-regulated convertase 1; Proprotein convertase 9 |
| *PLAT* | P00750 | *PLAT*; plasminogen activator, tissue type; plasminogen activator, tissue; T-PA; TPA; tissue-type plasminogen activator; plasminogen/activator kringle; reteplase; t-plasminogen activator |
| *POSTN* | Q15063 | *POSTN*; periostin; OSF-2; PN; osteoblast specific factor 2; periostin, osteoblast specific factor; OSF2; PDLPOSTN; osteoblast specific factor 2 (fasciclin I-like); periodontal ligament-specific periostin; Osteoblast-specific factor 2 |
| *PRKAR2B* | P31323 | *PRKAR2B*; Protein Kinase CAMP-Dependent Type II Regulatory Subunit Beta; Protein Kinase CAMP-Dependent Regulatory Subunit Type II Beta; CAMP-Dependent Protein Kinase Type II-Beta Regulatory Subunit; Protein Kinase CAMP-Dependent Regulatory Type II Beta; PRKAR2; CAMP-Dependent Protein Kinase Type II-Beta Regulatory Chain; WUGSC:H_RG363E19.2; H_RG363E19.2; RII-BETA |
| *PRKCA* | P17252 | *PRKCA*; protein kinase C alpha; PKC+¦; protein kinase C, alpha; AAG6; PKC-alpha; PKCA; PKCI+/-; PKCalpha; PRKACA; protein kinase C alpha type; PKC-A; aging-associated gene 6; PKC+Ä-¦ |
| *PRSS1* | P07477 | *PRSS1*; Serine Protease 1; TRY1; Cationic Trypsinogen; Protease Serine 1; Anionic Trypsin I; Pretrypsinogen I; Beta-Trypsin; EC 3.4.21.4; TRYP1; TRP1; Nonfunctional Trypsin 1; Digestive Zymogen; Anionic Trypsin-I; TCR V Beta 4.1; Trypsinogen 1; Trypsinogen A; Trypsin 1; Trypsin I; Trypsin-1; EC 3.4.21; TRY4 |
| *RAB24* | Q969Q5 | *RAB24*; RAB24; Member RAS Oncogene Family; Ras-Related Protein Rab-24 |
| *RHOT1* | Q8IXI2 | *RHOT1*; Ras Homolog Family Member T1; MIRO-1; MIRO1; ARHT1; Ras Homolog Gene Family Member T1; Mitochondrial Rho (MIRO) GTPase 1; Mitochondrial Rho GTPase 1; FLJ11040; Rac-GTP Binding Protein-Like Protein; Rac-GTP-Binding Protein-Like Protein; EC 3.6.5.-; EC 3.6.5; HMiro-1 |
| *SCGN* | O76038 | *SCGN*; secretagogin, EF-hand calcium binding protein; DJ501N12.8; SEGN; CALBL; calbindin-like; setagin; secretagogin |
| *SCPEP1* | Q9HB40 | *SCPEP1*; Serine Carboxypeptidase 1; RISC; Retinoid-Inducible Serine Carboxypeptidase; Serine Carboxypeptidase 1 Precursor Protein; Retinoid Inducible Serine Carboxypeptidase; EC 3.4.16.-; EC 3.4.16; HSCP1; SCP1 |
| *SERPINC1* | P01008 | *SERPINC1*; serpin family C member 1; ATIII; MGC22579; antithrombin III; signal peptide antithrombin part 1; coding sequence signal peptide antithrombin part 1; antithrombin (aa 375-432); serine (or cysteine) proteinase inhibitor, clade C (antithrombin), member 1; serpin peptidase inhibitor, clade C (antithrombin), member 1; AT3; AT3D; ATIII-R2; ATIII-T1; ATIII-T2; THPH7; serpin peptidase inhibitor clade C member 1; Antithrombin-III; Serpin C1 |
| *SERPINH1* | P50454 | *SERPINH1*; serpin family H member 1; HSP47; collagen binding protein 1; colligin; heat shock protein 47; serine (or cysteine) proteinase inhibitor, clade H (heat shock protein 47), member 2; serine (or cysteine) proteinase inhibitor, clade H (heat shock protein 47), member 1, (collagen binding protein 1); serpin peptidase inhibitor, clade H (heat shock protein 47), member 1, (collagen binding protein 1); AsTP3; CBP1; CBP2; OI10; PIG14; PPROM; RA-A47; SERPINH2; gp46; serpin H1; 47 kDa heat shock protein; arsenic-transactivated protein 3; cell proliferation-inducing gene 14 protein; colligin-1; colligin-2; rheumatoid arthritis antigen A-47; rheumatoid arthritis-related antigen RA-A47; serine (or cysteine) proteinase inhibitor, clade H (heat shock protein 47), member 2, (collagen-binding protein 2); Collagen-binding protein |
| *SFRP4* | Q6FHJ7 | *SFRP4*; secreted frizzled related protein 4; frpHE; FRP-4; FRZB-2; secreted frizzled-related protein 4; PYL; sFRP-4; frizzled protein, human endometrium; secreted frizzled-related protein 4; secreted frizzled-related protein 4 |
| *SHMT1* | P34896 | *SHMT1*; Serine Hydroxymethyltransferase 1; SHMT; CSHMT; Cytoplasmic Serine Hydroxymethyltransferase; Serine Hydroxymethyltransferase 1 (Soluble); Serine Hydroxymethyltransferase Cytosolic; Glycine Hydroxymethyltransferase; Serine Methylase; EC 2.1.2.1; MGC15229; MGC24556; 14 KDa Protein |
| *SMAD1* | Q15797 | *SMAD1*; SMAD family member 1; MADR1; JV4-1; MAD, mothers against decapentaplegic homolog 1 (Drosophila); SMAD, mothers against DPP homolog 1 (Drosophila); BSP-1; BSP1; JV41; MADH1; mothers against decapentaplegic homolog 1; MAD, mothers against decapentaplegic homolog 1; Mad-related protein 1; SMAD, mothers against DPP homolog 1; TGF-beta signaling protein 1; mothers against DPP homolog 1; transforming growth factor-beta signaling protein 1; MAD homolog 1; SMAD 1; hSMAD1; Transforming growth factor-beta-signaling protein 1 |
| *SOCS3* | O14543 | *SOCS3*; Suppressor Of Cytokine Signaling 3; SOCS-3; SSI-3; CIS3; Cytokine-Inducible SH2 Protein 3; STAT-Induced STAT Inhibitor 3; Cish3; SSI3; ATOD4; CISH3; CIS-3 |
| *SPART* | Q8N0X7 | *SPART*; spartin; KIAA0610; TAHCCP1; spastic paraplegia 20 (Troyer syndrome); SPG20; trans-activated by hepatitis C virus core protein 1; Spastic paraplegia 20 protein |
| *SPATA22* | Q8NHS9 | *SPATA22*; Spermatogenesis Associated 22; NYD-SP20; Spermatogenesis-Associated Protein 22; Testis Development Protein NYD-SP20; Testicular Tissue Protein Li 186; NYDSP20 |
| *STAT1* | P42224 | *STAT1*; signal transducer and activator of transcription 1; STAT91; ISGF-3; transcription factor ISGF-3 components p91/p84; signal transducer and activator of transcription 1, 91kD; signal transducer and activator of transcription 1, 91kDa; CANDF7; IMD31A; IMD31B; IMD31C; signal transducer and activator of transcription 1-alpha/beta |
| *STAT3* | P40763 | *STAT3*; signal transducer and activator of transcription 3; APRF; signal transducer and activator of transcription 3 (acute-phase response factor); ADMIO; ADMIO1; HIES; DNA-binding protein APRF; acute-phase response factor |
| *SYK* | P43405 | *SYK*; spleen associated tyrosine kinase; spleen tyrosine kinase; IMD82; p72-Syk; tyrosine-protein kinase SYK |
| *TBC1D13* | Q9NVG8 | *TBC1D13*; TBC1 Domain Family Member 13; FLJ10743; Epididymis Secretory Sperm Binding Protein |
| *TF* | P02787 | *TF*; transferrin; PRO1557; PRO2086; serotransferrin; HEL-S-71p; TFQTL1; beta-1 metal-binding globulin; epididymis secretory sperm binding protein Li 71p; siderophilin |
| *USP8* | P40818 | *USP8*; ubiquitin specific peptidase 8; HumORF8; KIAA0055; UBPY; SPG59; Ubiquitin carboxyl-terminal hydrolase 8; Ubiquitin isopeptidase Y; ubiquitin specific protease 8; PITA4; deubiquitinating enzyme 8; ubiquitin thiolesterase 8; ubiquitin-specific-processing protease 8; hUBPy; Ubiquitin thioesterase 8 |
| *WFIKKN2* | Q8TEU8 | *WFIKKN2*; WAP, follistatin/kazal, immunoglobulin, kunitz and netrin domain containing 2; WFIKKNRP; WFDC20B; WAP four-disulfide core domain 20B; GASP-1; hGASP-1; WAP, Kazal, immunoglobulin, Kunitz and NTR domain-containing protein 2; WAP, FS, Ig, two KU and NTR module related protein; WAP, follistatin, immunoglobulin, kunitz and NTR domain-containing-related protein; WFIKKN-related protein; growth and differentiation factor-associated serum protein 1; multivalent protease inhibitor protein; GASP1 |
| *YWHAB* | P31946 | *YWHAB*; Tyrosine 3-Monooxygenase/Tryptophan 5-Monooxygenase Activation Protein Beta; Tyrosine 3-Monooxygenase/Tryptophan 5-Monooxygenase Activation Protein Alpha Polypeptide; Tyrosine 3-Monooxygenase/Tryptophan 5-Monooxygenase Activation Protein Beta Polypeptide; 14-3-3 Protein Beta/Alpha; 14-3-3 Alpha; Protein 1054; KCIP-1; YWHAA; Protein Kinase C Inhibitor Protein-1; Protein Kinase C Inhibitor Protein 1; Epididymis Secretory Protein Li 1; 14-3-3 Beta; HEL-S-1; GW128; HS1 |
| *YWHAZ* | P63104 | *YWHAZ*; tyrosine 3-monooxygenase/tryptophan 5-monooxygenase activation protein zeta; KCIP-1; 14-3-3-zeta; 14-3-3 zeta; 14-3-3 delta; tyrosine 3-monooxygenase/tryptophan 5-monooxygenase activation protein, delta polypeptide; tyrosine 3-monooxygenase/tryptophan 5-monooxygenase activation protein, zeta polypeptide; HEL-S-3; HEL-S-93; HEL4; POPCHAS; YWHAD; 14-3-3 protein zeta/delta; 14-3-3 protein/cytosolic phospholipase A2; epididymis luminal protein 4; epididymis secretory protein Li 3; epididymis secretory protein Li 93; phospholipase A2; protein kinase C inhibitor protein-1; tyrosine 3/tryptophan 5 -monooxygenase activation protein, zeta polypeptide; Protein kinase C inhibitor protein 1 |

| **S7 Table. Enriched pathways in Mass Spec - SomaScan concordant genes** | | | | | |
| --- | --- | --- | --- | --- | --- |
| **Category** | **ID** | **Description** | **STR** | **FDR** | **Matching proteins in this network** |
| Reactome | HSA-381426 | Regulation of Insulin-like Growth Factor (IGF) transport and uptake by Insulin-like Growth Factor Binding Proteins (IGFBPs) | 2.02 | 3.24E-05 | PCSK9, SERPINC1, TF, KNG1 |
| Reactome | HSA-8957275 | Post-translational protein phosphorylation | 2.09 | 3.24E-05 | PCSK9, SERPINC1, TF, KNG1 |
